# The feedback regulator Nord controls Dpp/BMP signaling via extracellular interaction with Dally in the *Drosophila* wing

**DOI:** 10.1101/2021.03.11.434947

**Authors:** Takuya Akiyama, Chris W. Seidel, Matthew C. Gibson

**Affiliations:** Stowers Institute for Medical Research, Kansas City, MO 64110, USA; Department of Anatomy and Cell Biology, The University of Kansas School of Medicine, Kansas City, KS 66160, USA

**Keywords:** Nord, Dpp/BMP, Dally, NDNF, *Drosophila*, wing development

## Abstract

The *Drosophila* BMP 2/4 homologue Decapentaplegic (Dpp) acts as a morphogen to regulate diverse developmental processes, including wing morphogenesis. Transcriptional feedback regulation of this pathway ensures tightly controlled signaling outputs to generate the precise pattern of the adult wing. Nevertheless, few direct Dpp target genes have been explored and our understanding of feedback regulation remains incomplete. Here, we employ transcriptional profiling following *dpp* conditional knockout to identify *nord*, a novel Dpp/BMP feedback regulator. Nord mutants generated by CRISPR/Cas9 mutagenesis produce a smaller wing and display low penetrance venation defects. At the molecular level, *nord* encodes a heparin-binding protein and we show that its overexpression is sufficient to antagonize Dpp/BMP signaling. Further, we demonstrate that Nord physically and genetically interacts with the Dpp/BMP co-receptor Dally. In sum we propose that Nord acts with Dally to fine tune Dpp/BMP signaling, with implications for both developmental and disease models.

**Impact statement:** Functional analyses of the *Drosophila* homologue of Neuron Derived Neurotrophic Factor reveal a new mode of extracellular heparan sulfate proteoglycan regulation required for proper morphogen action.

## Introduction

For over 100 years, secreted factors have been recognized as powerful agents to control tissue growth and spatial patterning during development (Rogers and Schier, 2011). This idea gained particular experimental traction from Spemann and Mangold’s organizer transplantation experiment (Spemann and Mangold, 1924). When the dorsal blastopore lip of a weakly pigmented salamander embryo is grafted to the ventral side of a pigmented host, the grafted fragment induces secondary axis formation in neighboring pigmented cells, causing a conjoined twin. Deciphering the chemical basis for this organizer activity, however, remained elusive. In the 1990s, a fuller and more complex picture of cell-cell signaling began to emerge. Molecular analyses of the organizer identified many secreted antagonists of signaling molecules, such as Chordin for Bone morphogenetic protein (BMP), Frizzled-related Protein 1 for Wnt and Cerberus for Nodal (Sasai et al., 1994; Sasai et al., 1995; Bouwmeester et al., 1996; Leyns et al., 1997). These secreted signaling molecules are repeatedly utilized during development and homeostasis to establish proper cell-cell communication and their misregulation often leads to congenital abnormalities and diseases (Wu et al., 2016; Li and Elowitz, 2019; Zhang and Wang, 2020).

During development, a wide range of signaling molecules act as morphogens to govern diverse processes, such as embryonic dorsoventral (D/V) axis formation and neural and limb patterning in both vertebrates and invertebrates (O’Connor et al., 2006; Affolter and Basler, 2007; Muller et al., 2013; Akiyama and Gibson, 2015b; Bier and De Robertis, 2015). Morphogens provide spatial information to cells in a morphogenetic field by creating a concentration gradient, and direct distinct cell fates in a concentration-dependent manner. Over the last three decades, analysis of the developing *Drosophila* wing has greatly contributed to our understanding the molecular basis of morphogen action (Lecuit et al., 1996; Nellen et al., 1996; O’Connor et al., 2006; Affolter and Basler, 2007; Akiyama and Gibson, 2015b; Beira and Paro, 2016). The adult wing develops from a primordium called the wing disc, an epithelial invagination which grows dramatically during larval stages and then everts during metamorphosis to produce the adult structure. During wing development, Wingless (Wg; a *Drosophila* Wnt), Hedgehog (Hh) and Decapentaplegic (Dpp; a *Drosophila* BMP2/4) act as morphogens to pattern the wing (O’Connor et al., 2006; Affolter and Basler, 2007; Gradilla and Guerrero, 2013; Akiyama and Gibson, 2015b; Beira and Paro, 2016; Bejsovec, 2018). Among them, Dpp is perhaps the most studied morphogen and exhibits two distinct morphogen actions during this process. In the wing disc, Dpp emanates from a stripe of cells adjacent to the anterior-posterior (A/P) boundary to generate a long-range Dpp morphogen gradient along the A/P axis (Lecuit et al., 1996; Nellen et al., 1996; Entchev et al., 2000; Teleman and Cohen, 2000). Dpp activates downstream signaling through a tetrameric receptor complex consisting of BMP type I and type II receptors (Brummel et al., 1994; Nellen et al., 1994; Penton et al., 1994; Xie et al., 1994; Letsou et al., 1995; Ruberte et al., 1995). In *Drosophila*, the BMP type I receptors, such as Thickveins (Tkv), phosphorylate the downstream signal transducer Mothers against dpp (Mad), resulting in a phosphorylated Mad (p-Mad) gradient that provides a direct readout of Dpp/BMP signaling activity across the wing disc (Newfeld et al., 1996; Tanimoto et al., 2000). The graded distribution of Dpp/BMP activity in turn induces the nested expression of target genes, including *Daughters against dpp* (*Dad*)*, spalt* (*sal*), *optomotor blind* (*omb*) and *brinker* (*brk*), to establish the intervein and longitudinal vein regions. During pupal development, Dpp/BMP signaling is further required for wing vein morphogenesis (O’Connor et al., 2006; Blair, 2007; Raftery and Umulis, 2012). The Dpp morphogen, now expressed in the longitudinal veins, not only activates the signaling pathway locally but is also transported into the presumptive crossvein region to shape the morphogen gradient by the actions of extracellular BMP interacting proteins, such as Short gastrulation (Sog, a *Drosophila* Chordin) (Conley et al., 2000; Ralston and Blair, 2005; Serpe et al., 2005; Shimmi et al., 2005). Dpp/BMP activation in the longitudinal vein and crossvein regions triggers the terminal wing vein differentiation process.

Transcriptional feedback regulation of Dpp/BMP signaling is an essential mechanism for buffering against genetic and environmental perturbations (O’Connor et al., 2006; Lander, 2011; Raftery and Umulis, 2012; Bier and De Robertis, 2015). Previous studies have discovered several Dpp/BMP feedback regulators, such as Division abnormally delayed (Dally), Pentagone (Pent) and Larval translucida (Ltl) (Fujise et al., 2003; Vuilleumier et al., 2010; Szuperak et al., 2011). Dally is a member of the glypican family of heparan sulfate proteoglycans (HSPGs), which consist of a protein core and covalently attached heparan sulfate glycosaminoglycan (HS) chains. Dally localizes on the cell membrane through a glycosylphosphatidylinositol anchor and serves as a major Dpp/BMP co-receptor (Dejima et al., 2011; Nakato and Li, 2016). Dally forms a complex with Dpp and stabilizes it in the extracellular space, thus facilitating the formation of the Dpp morphogen gradient (Akiyama et al., 2008). A recent study demonstrates that Pent internalizes Dally and Dally-like, another *Drosophila* glypican, from the cell surface via a Rab5-dependent endocytosis, and this Pent-mediated degradation of glypicans plays a critical role in gradient formation (Norman et al., 2016). In addition, another extracellular Dpp/BMP feedback regulator, Ltl, physically interacts with Dally-like and antagonizes Dpp/BMP signaling in both the wing disc and the pupal wing (Szuperak et al., 2011). Glypican activity is thus a key factor in orchestrating Dpp morphogen actions in the developing wing. In this study, we use a novel conditional mutagenesis approach to elucidate targets of Dpp signaling in the wing disc and identify the secreted Dpp/BMP feedback regulator Nord, a *Drosophila* Neuron-Derived Neurotrophic Factor (NDNF) homologue. *nord* mutants develop smaller wings and exhibit wing venation defects at low penetrance. Further consistent with a role in feedback regulation, overexpression of Nord antagonizes Dpp/BMP signaling. In addition, we show that Nord binds to Dally and that loss of *nord* suppresses the wing vein defects normally observed in *dally* mutants. Based on these data, we propose a model in which Nord functions with Dally to modulate Dpp/BMP signaling during wing development.

## Results

### Dpp/BMP target genes in the developing wing disc

Due to a lack of techniques that completely disrupt *dpp* function, transcriptional targets of Dpp/BMP in the developing wing have been identified by relatively indirect approaches, such as genome-wide *in silico* screening of Dpp/BMP silencer elements and microarray comparison between *wild-type* cells and BMP type I receptor *tkv* mutant clones (Vuilleumier et al., 2010; Szuperak et al., 2011; Organista et al., 2015). Recently, advances in genome editing technology have led to the creation of three *dpp* conditional knockout lines, *dpp^FO^*, *dpp^CA^* and *dpp^PSB^* (Akiyama and Gibson, 2015a; Bosch et al., 2017). Using *flippase recombination target sequence* (*FRT*)-mediated excision of the endogenous *dpp* locus, these lines allow for conditional elimination of *dpp* function in the wing disc and thus an opportunity to elucidate Dpp/BMP target genes using mRNA sequencing analysis.

Hypothetically, animals homozygous for conditionally inactivatable alleles should feature fully *wild-type* activity before *FRT*-mediated excision and null activity after. In order to identify the most suitable *dpp* conditional allele for RNA-Seq profiling, we first examined homozygous animals for *dpp* phenotypes and tested each line using conventional complementation testing (Figure 1A and B). In the absence of conditional inactivation, we found that a majority of *dpp^PSB^* homozygotes (63%) displayed a posterior crossvein (PCV) phenotype, which was observed in just 3.8% of *dpp^CA^* adults and was absent in *dpp^FO^* flies (Figure 1A). Since PCV development is sensitive to Dpp/BMP activity (Conley et al., 2000; Ralston and Blair, 2005; Serpe et al., 2005; Shimmi et al., 2005), this suggests that the genome editing of *dpp^PSB^* may mildly interfere with *dpp* function. *dpp* is known as a haploinsufficient gene, and its gene dosage is critical for viability (Irish and Gelbart, 1987; Wharton et al., 1996). Thus, to directly assess *dpp* activity in all three conditional knockout lines, we also performed an adult viability assay by crossing these conditional lines to weak (*dpp^hr56^*) and strong (*dpp^hr27^*) loss of function alleles (Figure 1B) (Spencer et al., 1982; Irish and Gelbart, 1987; Wharton et al., 1996). Consistent with loss of *dpp* function in the absence of *FRT*-mediated excision, *dpp^PSB^*/*dpp^hr56^* and *dpp^PSB^/dpp^hr27^* flies showed reduced adult viability (26.3% and 13.6% survival, respectively; Figure 1B). Unexpectedly, *dpp^CA^*, whose homozygotes rarely exhibited PCV defects, had the worst survival outcomes. Only 4% of *dpp^CA^*/*dpp^hr56^* animals survived to adulthood, and *dpp^CA^* completely failed to complement *dpp^hr27^* (Figure 1B). These results indicate significant loss of function prior to conditional inactivation. In contrast, *dpp^FO^* had the least impact on *dpp* activity. 40.1% of *dpp^FO^*/*dpp^hr27^* animals survived to adulthood and *dpp^FO^*/*dpp^hr56^* showed a similar viability to *+*/*dpp^hr56^* controls (85.1%; Figure 1B). We therefore used the *dpp^FO^* line for further analysis.

**Figure 1:**
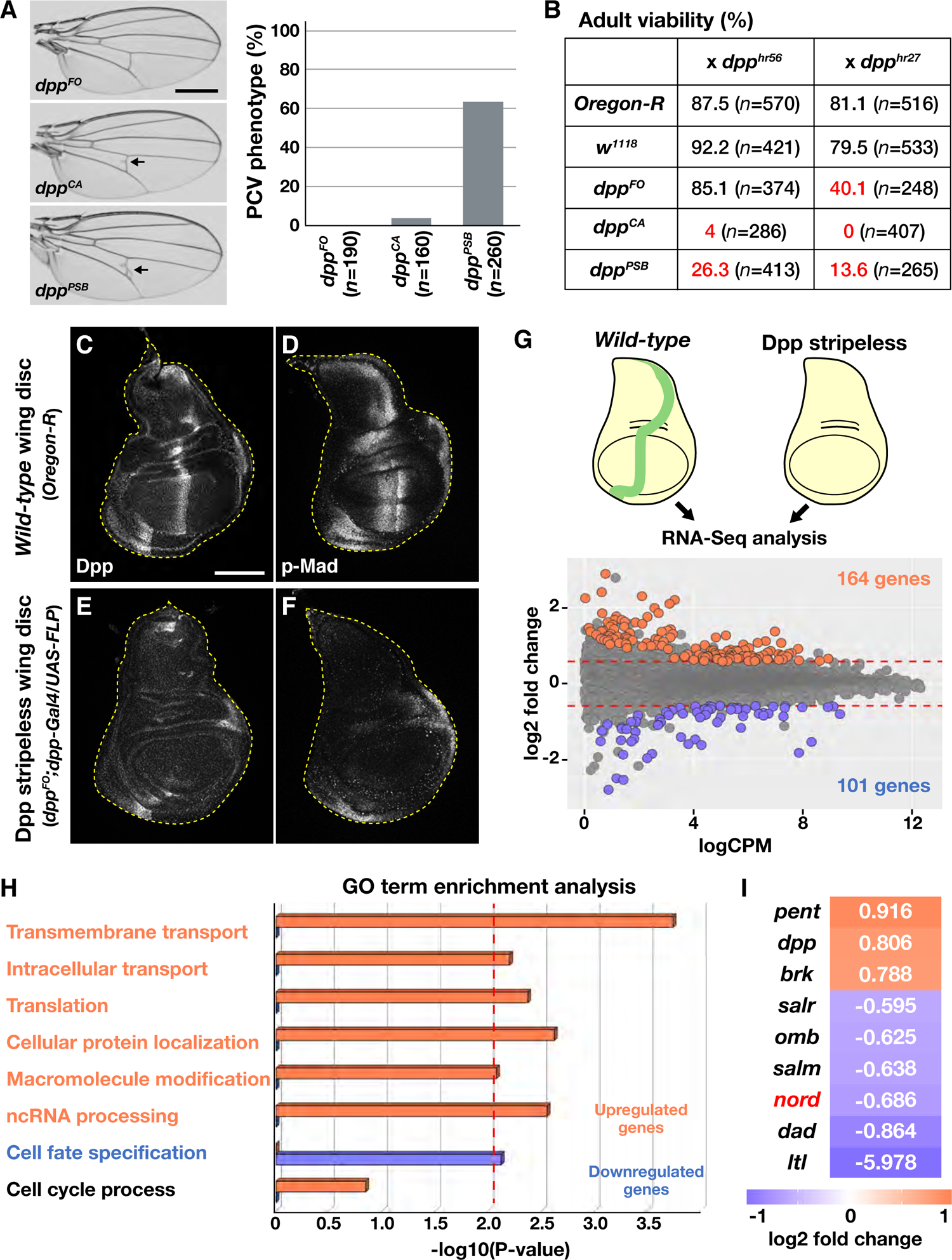
Identification of Dpp/BMP target genes. (**A**, **B**) Characterization of three *dpp* conditional knockout lines. (**A**) Adult wings from *dpp* conditional knockout homozygous animals. Two lines, *dpp^CA^* and *dpp^PSB^*, exhibited the PCV spur phenotype, a typical PCV defect associated with aberrant Dpp/BMP signaling activity. *dpp^FO^* (*n*=190). *dpp^CA^* (*n*=160). *dpp^PSB^* (*n*=219). Scale bar: 0.5 mm. Anterior is oriented to the top side of images. (**B**) Adult viability of transheterozygotes carrying a *dpp* conditional knockout chromosome and a *dpp* hypomorphic allele. Two classical hypomorphic alleles, a weak *dpp^hr56^* and strong *dpp^hr27^*, were used for the crosses. Several combinations of transheterozygotes showed low viabilities (*red*). (**C**-**F**) Representative images of Dpp expression and Dpp/BMP activity (p-Mad) gradient in *wild-type* and Dpp stripeless wing discs. Scale bar: 100 μm. Anterior is to the left of all figures. (**G**) Transcriptome comparison between *wild-type* and Dpp stripeless wing discs. Significantly up and downregulated genes are highlighted. (**H**) Gene ontology (GO) term enrichment analysis of up and downregulated genes in Dpp stripeless wing disc. *Red* dashed line indicates an adjusted *P*-value cutoff of 0.01. (**I**) RNA-Seq analysis revealed well-known Dpp/BMP target genes and a novel target gene, *nord*.

As reported in previous studies, the endogenous stripe of Dpp expression is consistently eliminated in *dpp^FO^/dpp^FO^; dpp-Gal4/UAS-FLP* (Dpp stripeless) animals via flippase-mediated recombination. This, in turn, results in loss of the Dpp/BMP activity (p-Mad) gradient in third-instar wing discs (Figure 1C-F) (Akiyama and Gibson, 2015a). We used this approach to discover Dpp/BMP target genes by comparing the transcriptomes of *wild-type* and Dpp stripeless wing discs, identifying 164 genes upregulated in the absence of *dpp* and 101 that were downregulated (an adjusted *P*-value < 0.01 and a fold change > 1.5; Figure 1G; supplemental excel file). Gene ontology (GO) term enrichment analysis revealed that genes regulating cell fate specification were enriched in the downregulated gene cluster, whereas upregulated genes were involved in biological processes such as transmembrane transport and cellular protein localization (Figure 1H). Further, consistent with the mild growth phenotype of Dpp stripeless wing discs, cell cycle associated genes were not significantly enriched in either population (Figure 1H). Validating our experimental approach, transcriptional profiling identified multiple well-known Dpp/BMP target genes, including the feedback regulators *Dad*, *pent* and *ltl*, (Figure 1I) (Nellen et al., 1996; Tsuneizumi et al., 1997; Campbell and Tomlinson, 1999; Minami et al., 1999; Vuilleumier et al., 2010; Szuperak et al., 2011). In addition to known Dpp/BMP targets, we identified a novel Dpp/BMP target gene, *nord* (Figure 1I).

### Dpp/BMP regulates *nord* expression in the wing disc

We performed RNA fluorescent *in situ* hybridization (FISH) to visualize *nord* expression in wing discs and observed robust expression throughout the third-instar larval stage (Figure 2A-F; Figure 2-figure supplement 1A-H). Similar to *dpp*, *nord* was strongly expressed in a stripe of cells at the A/P compartment boundary. Interestingly, *nord* expression was repressed along the D/V boundary (Figure 2A-F; Figure 2-figure supplement 1A-E). Especially strong *nord* mRNA signal was detected in the epithelial folds located in the central part of wing pouch (Figure 2-figure supplement 1F-H). Intriguingly, unlike other well-characterized Dpp/BMP target genes, *nord* was exclusively expressed in anterior cells (Figure 2A-F; Figure 2-figure supplement 1A-E) (Nellen et al., 1996; Tsuneizumi et al., 1997; Campbell and Tomlinson, 1999; Minami et al., 1999; Vuilleumier et al., 2010; Szuperak et al., 2011). Further, in the pupal wing 24-hours after puparium formation (APF), *nord* mRNA was only detected in a row of distal anterior cells where neither Dpp transcription nor Dpp/BMP activity is observed (Figure 2-figure supplement 1I and J). These results raise the possibility that *nord* expression is not solely regulated by Dpp/BMP signaling during development.

**Figure 2:**
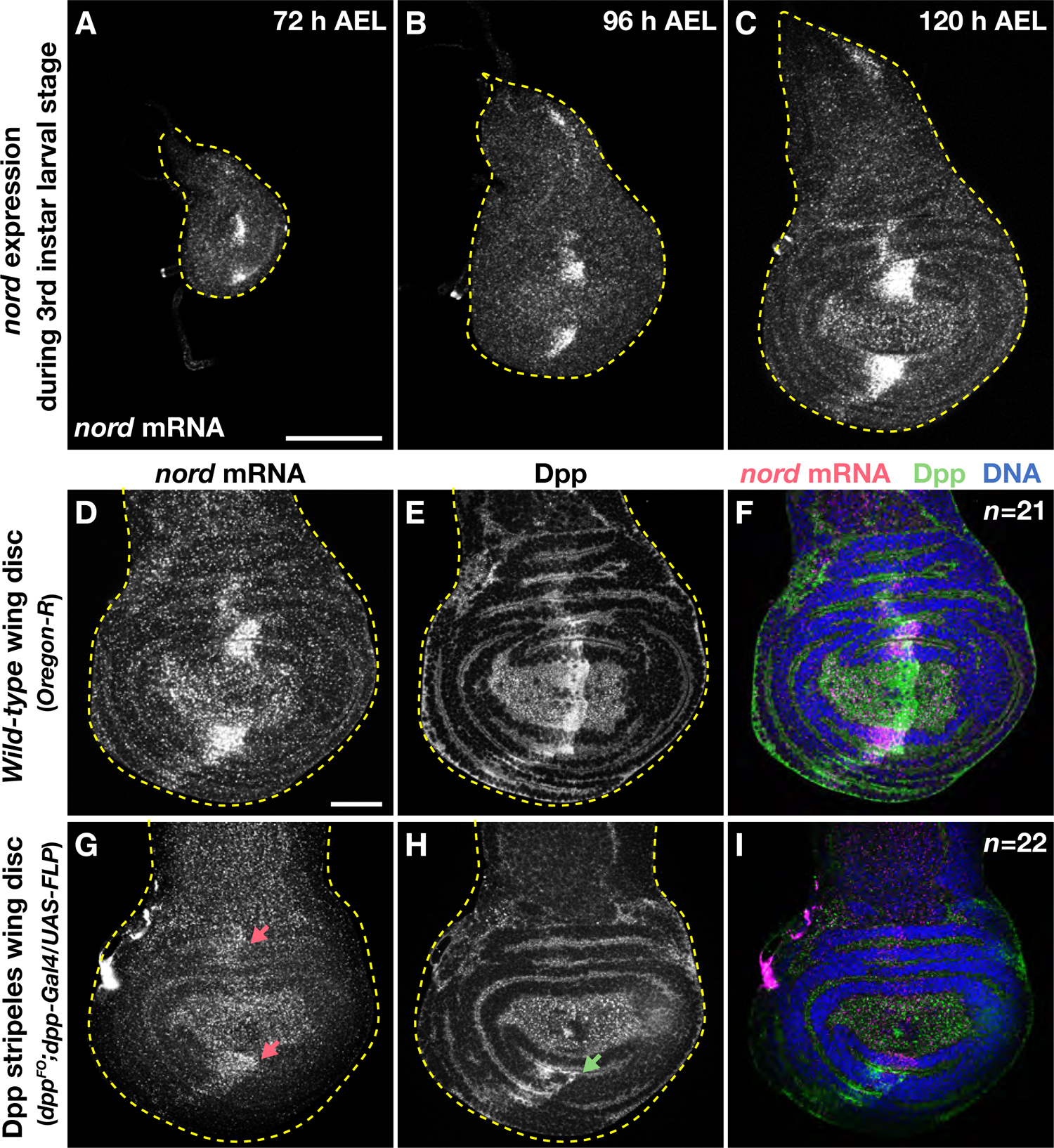
Dpp regulates *nord* expression in the wing disc. (**A**-**C**) *nord* expression patterns in the third-instar wing discs (72-120 h AEL). (**D**-**I**) *nord* RNA FISH and Dpp staining in *wild-type* and Dpp stripeless wing discs. *nord* expression was dramatically reduced in Dpp stripeless wing disc (*magenta* arrows). A *green* arrow indicates a residual Dpp expression in Dpp stripeless wing disc. Scale bars: 100 μm for **A**, 50 μm for **D**. Anterior is oriented to the left of all panels.

We next tested whether *dpp* was required for *nord* expression (Figure 2D-I). Consistent with our RNA-Seq data, eliminating the stripe Dpp from the wing disc led to a strong reduction of *nord* expression (Figure 2G-I). To further investigate the requirement of Dpp/BMP signaling, we performed clonal analyses using the FLP-OUT technique to ectopically inhibit or activate the Dpp/BMP signaling pathway (Figure 2-figure supplement 2A-F) (Ito et al., 1997). As expected, we found that ectopic inhibition of Dpp/BMP signaling by Dad overexpression significantly reduced *nord* expression (Figure 2-figure supplement 2C and D). Surprisingly, cell autonomous activation of Dpp/BMP signaling via Tkv^QD^ overexpression did not induce *nord* (Figure 2-figure supplement 2E and F). Since *nord* mRNA levels were low at the D/V boundary where Wg expression is high, we also examined whether ectopic Wg/Wnt pathway activation is sufficient to repress *nord*. We generated *arm^S10^* FLP-OUT clones throughout the wing disc to activate Wg/Wnt signaling but found that *nord* expression was not affected (Figure 2-figure supplement 2G and H). Collectively, these results suggest that *nord* expression is not downstream of Wg/Wnt signaling and that Dpp/BMP signaling is required but not sufficient to drive *nord* expression in the wing disc.

### *nord* governs growth and patterning during wing development

To determine if *nord* gene function is necessary for wing development, we used CRISPR/Cas9 mutagenesis to generate two deletion alleles, *nord^Δ1728^* and *nord^Δ1^* (Figure 3A; Figure 3-figure supplement 1). *nord* encodes a secreted protein with an NDNF motif, which is essential for its function (Figure 3A). Both deletions caused frameshifts, resulting in predicted protein truncations lacking large portions of the NDNF domain (Figure 3A; Figure 3-figure supplement 1). Since the *nord^Δ1728^* deletion resulted in an earlier termination codon compared to the *nord^Δ1^* deletion, we mainly used this mutant allele for subsequent functional analyses. To determine how loss of *nord* affects Dpp/BMP signaling during wing development, we stained *nord^Δ1728^* homozygous mutant discs with anti-p-Mad antibodies (Figure 3B-D). We found that both the control and *nord^Δ1728^* discs show similar p-Mad gradients, though the mutant discs had a slightly higher anterior p-Mad peak (Figure 3B-D). Second, we checked Dpp/BMP activity in the 24-hour APF pupal wing. At this stage, p-Mad signal was detected in both future longitudinal veins and the crossvein regions (Figure 3E and G). Although the majority of mutant pupal wings showed comparable Dpp/BMP activity to controls, they did exhibit ectopic Dpp/BMP activity in the PCV region at low frequency (13%; Figure 3F and H). Finally, we investigated adult wing phenotypes (Figure 3I-L). As might be expected based on the presence of ectopic Mad phosphorylation, *nord^Δ1728^* homozygotes infrequently displayed a spur of ectopic venation in the PCV region (8.6%; Figure 3I-K). Importantly, we also found that *nord^Δ1728^* animals developed significantly smaller wings than the control (Figure 3I, J and L). Complementation tests with a genomic deficiency confirmed these findings: *nord^Δ1728^*/*Df(2R)BSC356* and *nord^Δ1^* homozygous animals show a similar PCV phenotype to *nord^Δ1728^* homozygotes (Figure 3K). These results rule out the possibility of off-target effects caused by CRISPR/Cas9-mediated mutagenesis and suggest that both *nord* mutants are likely null alleles.

**Figure 3:**
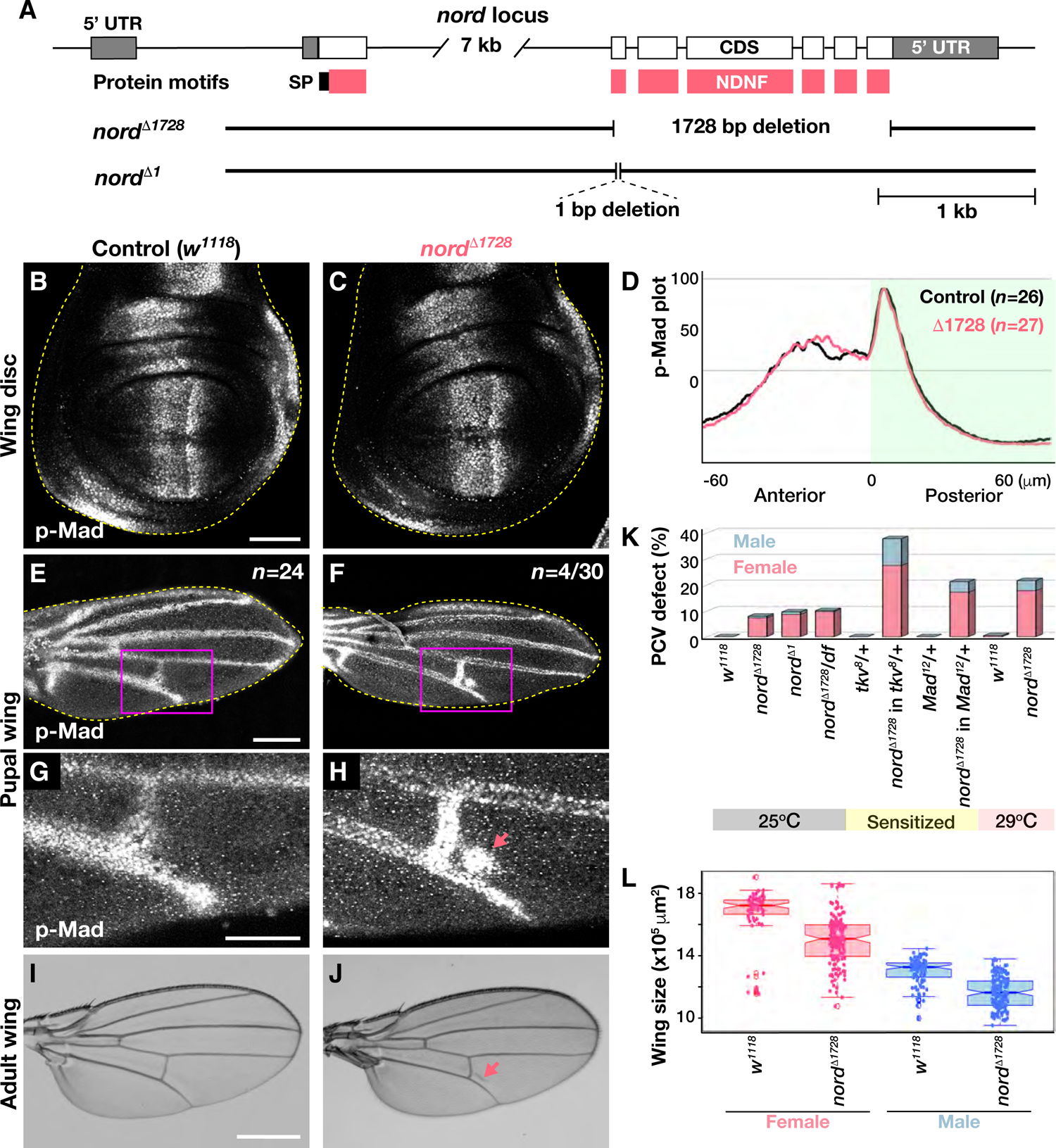
*nord* mutants exhibit wing growth and mild wing vein phenotypes. (**A**) The *nord* locus. Nord protein motifs and the lesions of the deletion mutants are indicated on the diagram. SP: signal peptide. NDNF: neuron-derived neurotrophic factor. (**B**, **C**) p-Mad gradients in control and *nord^Δ^*^1728^ mutant wing discs. (**D**) Averaged p-Mad intensity plot profiles. Control and *nord^Δ^*^1728^ mutant wing discs showed similar p-Mad intensity plot profiles. 0 represents the anterior-posterior compartment boundary. (**E-H**) p-Mad staining in 24-hour APF control and *nord^Δ1728^* mutant pupal wings. **G** and **H** show magnified images of the boxed regions in **E** and **F**, respectively. An ectopic Dpp/BMP activation was rarely observed in *nord^Δ1728^* mutant pupal wing (arrow). (**I**, **J**) Control and *nord^Δ1728^* mutant adult wings. *nord^Δ1728^* animals occasionally exhibited the PCV spur phenotype (arrow). **(K**) Quantification analysis of PCV phenotype in standard (25°C), sensitized genetic (*tkv^8^* and *Mad^12^* heterozygous backgrounds) and stressed (29°C) conditions. *w^1118^* (female at 25°C: *n*=70, male at 25°C: *n*=99; female at 29°C: *n*=120, male at 29°C: *n*=111). *nord^Δ1728^* (female at 25°C: *n*=151, male at 25°C: *n*=163; female at 29°C: *n*=132, male at 29°C: *n*=120). *nord^Δ1^* (female: *n*=110, male: *n*=104). *nord^Δ1728^/df* (female: *n*=123, male: *n*=133). *tkv^8^/+* (female: *n*=101, male: *n*=63). *tkv^8^, nord^Δ1728^/nord^Δ1728^* (female: *n*=115, male: *n*=82). *Mad^12^ /+* (female: *n*=82, male: *n*=80). *Mad^12^, nord^Δ1728^/nord^Δ1728^* (female: *n*=90, male: *n*=64). Intriguingly, females more frequently displayed the PCV spur than males. (**L**) Wing size. *nord^Δ1728^* wings were significantly smaller than control wings (*P*-value < 0.001). *w^1118^* (female: *n*=66, male: *n*=96). *nord^Δ1728^* (female: *n*=150; male: *n*=163). Scale bars: 50 μm for **B** and **G**, 100 μm for **E**, 0.5 mm for **I**. Anterior is oriented to the left in wing disc figures and to the top in pupal and adult wing images.

BMP feedback regulation plays a critical role in buffering against both genetic and environmental perturbations (O’Connor et al., 2006; Lander, 2011; Raftery and Umulis, 2012; Bier and De Robertis, 2015). Therefore, we tested if *nord* could function to buffer Dpp/BMP signaling after genetic perturbations or when animals were kept under environmentally stressful conditions. Indeed, the PCV phenotype caused by a loss of *nord* was enhanced in both *tkv* (a *Drosophila* BMP type I receptor) and *Mad* (a *Drosophila* Smad1/5/8) heterozygous mutant backgrounds (Figure 3K). Additionally, when *nord* mutant animals were reared at high temperature (29°C), the penetrance of the PCV defect was significantly increased (Figure 3K). Based on these results combined with Nord’s protein structure and its highly localized expression pattern during wing development, we propose that Nord acts as a secreted Dpp/BMP feedback regulator which functions to ensure robust developmental outcomes under sensitized genetic and stress conditions.

### *nord* antagonizes Dpp/BMP signaling

To gain a better understanding of how Nord functions in Dpp/BMP signaling, we overexpressed *nord* using the Gal4/*UAS* system (Figure 4) (Brand and Perrimon, 1993). *nord* overexpression in the wing pouch domain using *nub-Gal4* resulted in a reduced p-Mad gradient with a weaker posterior peak compared to controls (Figure 4A-C). *nord* overexpression caused a variety of wing defects, including reduced size and a consistent loss of the anterior cross vein (ACV; Figure 4D-G). We also observed a PCV defect and wing notching at a lower penetrance (Figure 4D-F). Although the crossvein defects are most likely caused by aberrant Dpp/BMP signaling activity, wing notching is linked to other pathways, including the Notch and Wg/Wnt signaling pathways (Zacharioudaki and Bray, 2014; Bejsovec, 2018). Therefore, we examined whether *nord* overexpression also affects Wg/Wnt signaling (Figure 4-figure supplement 1). As a Wg/Wnt activity reporter, fz3-RFP exhibited a graded signal centered at the D/V compartment boundary in the wing disc (Figure 4-figure supplement 1A and C) (Olson et al., 2011). However, when *nord* was overexpressed in the wing pouch, the amplitude of the Wg/Wnt activity gradient was significantly reduced (Figure 4-figure supplement 1A-C). Altogether, our results demonstrate that *nord* overexpression not only antagonizes Dpp/BMP signaling but also inhibits the Wg/Wnt signaling pathway.

**Figure 4:**
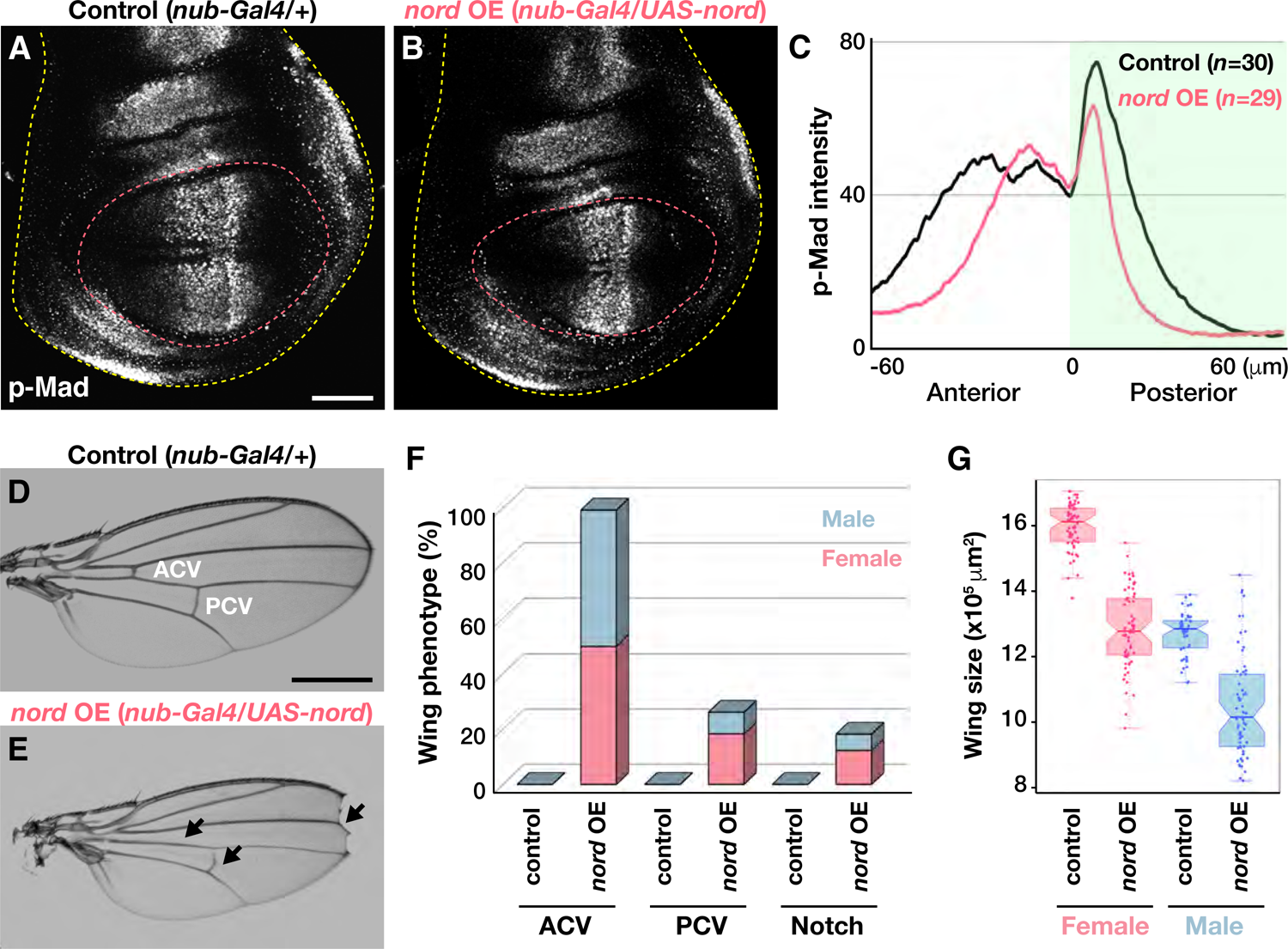
*nord* overexpression perturbs Dpp/BMP signaling (**A**, **B**) p-Mad gradients in control wing disc and the wing disc overexpressing *nord*. *nord* is overexpressed in the wing pouch region using *nub-Gal4* (*magenta* dashed line). (**C**) Averaged p-Mad intensity plot profiles. *nord* overexpression caused a narrower p-Mad gradient and a weaker posterior peak than control. The posterior compartment is shaded. (**D**, **E**) Adult wing phenotypes of control and *nord* overexpressing animals. *nord* overexpression in the wing pouch resulted in the ACV, PCV and notch phenotypes (arrows). (**F**) Quantitative analysis of adult wing phenotypes. (**G**) Wing size comparison. *nord* overexpression significantly reduced wing size (*P*-value < 0.001). Control (female: *n*=60, male: *n*=46). *nord* overexpression (female: *n*=64, male: *n*=62). Scale bars: 50 μm for **A**, 0.5 mm for **D**. Anterior is oriented to the left in wing disc figures and to the top in adult wing images.

### *nord* acts with HSPGs to modulate Dpp/BMP signaling

How does Nord regulate these signaling pathways in the wing disc? As mentioned above, *nord* encodes a secreted NDNF family protein predicted to possess heparin-binding activity in vertebrates (e.g. human; https://www.uniprot.org/uniprot/Q8TB73). Heparin is a variant of HS polysaccharides, which are ubiquitously present in all tissue types as a major component of HSPGs (Hammond et al., 2014; Nakato and Li, 2016; Xie and Li, 2019; Jayatilleke and Hulett, 2020). In the developing wing disc, Dally and Dally-like, members of the glypican family of HSPGs, regulate both the Dpp/BMP and Wg/Wnt signaling pathways by acting as co-receptors (Nakato and Li, 2016; Mii and Takada, 2020). Thus, we hypothesized that Nord acts with HSPGs to regulate these signaling pathways. To test this, we first examined subcellular localization and heparin binding activity of Nord using *Drosophila* S2 tissue culture cells (Figure 5A and B). S2 cells were transfected with *nord-HA* DNA to transiently express Nord-HA protein. We found that the signal peptide of Nord-HA protein was efficiently cleaved, and the mature Nord-HA protein was secreted into the conditioned media (Figure 5A). As expected, the secreted Nord-HA protein also bound to heparin (Figure 5A and B). Next, to test a physical interaction between Nord and Dally, a major co-receptor for Dpp/BMP signaling (Dejima et al., 2011), we performed co-immunoprecipitation experiments using S2 cell lysates containing Nord-HA with or without Dally-Myc. Dally-Myc protein was only recovered when S2 cells were co-transfected with both *nord-HA* and *dally-Myc* DNAs (Figure 5C). Further, we found that this physical interaction was not totally dependent on HS because Nord also binds to Dally^ΔGAG^, a mutant form of Dally lacking an HS modification (Figure 5C) (Kirkpatrick et al., 2006). Nevertheless, it seems likely that the presence of HS is able to enhance the physical interaction between Nord and Dally (Figure 5C).

**Figure 5:**
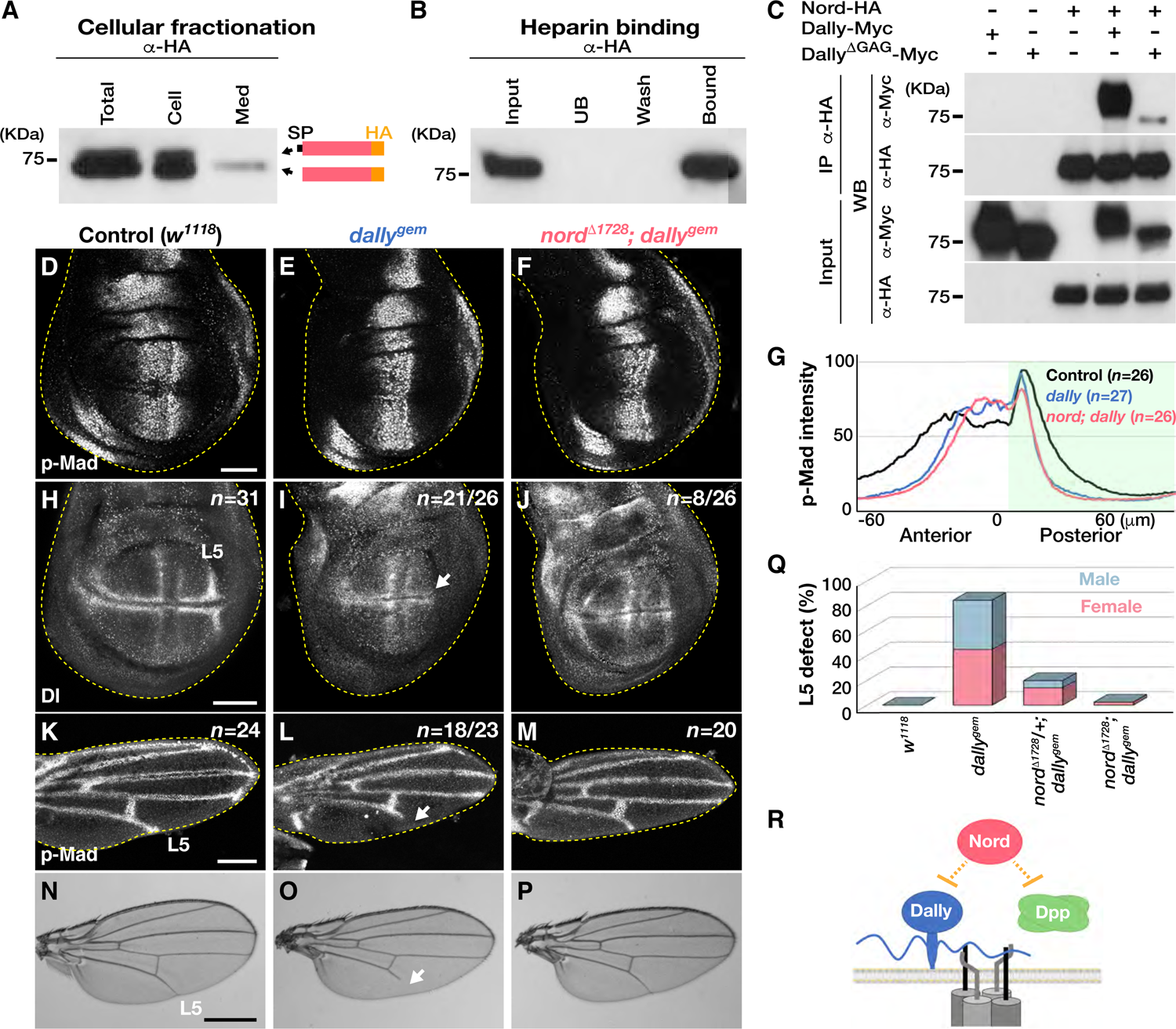
*nord* mutations suppress the L5 wing vein defect of the *dally* mutant. (**A**) Subcellular fractionation of S2 cells expressing Nord-HA. The cells were fractionated into total, cell and conditioned medium (Med) fractions. Nord-HA was detected in the conditioned medium fraction. SP: signal peptide. (**B**) Heparin-binding assay using conditioned media containing Nord-HA protein. Nord-HA proteins bound to heparin. UB: unbound. (**C**) Physical interaction between Nord-HA and Dally-Myc. S2 cell lysates containing Nord-HA with or without Dally-Myc or Dally^ΔGAG^-Myc, which lacks HS modifications, were used for the co-immunoprecipitation experiment. While both forms of Dally proteins were recovered with immunoprecipitated Nord-HA, Dally-Myc showed a stronger band than Dally^ΔGAG^-Myc. (**D**-**F**) Anti-p-Mad staining of control, *dally^gem^* and *nord^Δ1728^*; *dally^gem^* wing discs. (**G**) p-Mad intensity plot profiles. Similar p-Mad intensity plots were obtained from *dally^gem^* and *nord^Δ1728^*; *dally^gem^* wing discs. (**H**-**J**) Visualization of longitudinal wing vein primordial cells by anti-Dl staining. (**K**-**M**) Dpp/BMP activity (p-Mad) in 24-hour APF pupal wings. (**N**-**P**) Adult wing phenotypes. *dally^gem^* wing discs frequently lack L5 Dl staining (arrow in **I**) and show the L5 defect in pupal and adult wings (arrows in **L** and **O**). These phenotypes were significantly rescued in the double mutant (**J**, **M**, and **P**). Scale bars: 50 μm for **D** and **H**, 100 μm for **K**, 0.5 mm for **N**. Anterior is oriented to the left in wing disc panels, and to the top in pupal and adult wing figures. (**Q**) Quantification of the L5 vein defect. *w^1118^* (female: *n*=67, male: *n*=98). *dally^gem^* (female: *n*=160, male: *n*=117). *nord^Δ1728^/+; dally^gem^* (female: *n*=159, male: *n*=80). *nord^Δ1728^; dally^gem^* (female: *n*=146, male: *n*=76). (**R**) A model for Nord functions in Dpp/BMP signaling.

To examine how Nord affects Dally functions *in vivo*, we performed genetic interaction experiments (Figure 5D-Q; Figure 5-figure supplement 1A-C). Consistent with previous reports, *dally^gem^* mutants exhibited a sharp p-Mad gradient, lost the specification of L5 primordial cells in the wing disc, and developed a smaller wing lacking a distal portion of the L5 vein (Figure 5D, E, H, I, K, L, N and O) (Fujise et al., 2003; Akiyama et al., 2008). Although *nord^Δ1728^*; *dally^gem^* double mutant wing discs had similar p-Mad gradients to those observed in *dally^gem^* homozygotes, we observed that the majority of double mutant discs were able to specify the L5 vein primordia as visualized with anti-Delta (Dl) staining (Figure 5D-J). As a result, the double mutants developed an intact L5 vein in both pupal and adult wings (Figure 5K-Q). We also noticed that heterozygosity for *nord^Δ1728^* was sufficient to partially rescue the L5 defect, suggesting a strong genetic interaction (Figure 5Q). Further, the *nord* PCV defect was completely suppressed in *nord^Δ1728^*; *dally^gem^* double mutants, although wing notching associated with loss of *dally* was enhanced and the wing size defects were not rescued in the double mutant animals (Figure 5-figure supplement 1A-C). We therefore infer that loss of *nord* function both suppresses or enhances *dally* mutant phenotypes in a highly context-dependent manner. Altogether, these data suggest that Nord functions with the co-receptor Dally to modulate both Dpp/BMP and Wg/Wnt signaling.

## Discussion

### The Dpp/BMP morphogen gradient and wing disc growth

While the Dpp/BMP morphogen activity gradient is clearly essential for wing patterning, the mechanistic basis for its requirement in wing disc growth remains poorly understood (Akiyama and Gibson, 2015a; Barrio and Milan, 2017; Bosch et al., 2017; Matsuda and Affolter, 2017). Here, we found that three independently constructed *dpp* conditional knockout lines displayed distinct growth phenotypes when the Dpp stripe expression was disrupted (Akiyama and Gibson, 2015a; Bosch et al., 2017; Matsuda and Affolter, 2017). *dpp^FO^* caused mild growth defects, whereas two additional lines, *dpp^CA^* and *dpp^PSB^*, showed severe growth impairments (Bischof et al., 2007; Akiyama and Gibson, 2015a; Matsuda and Affolter, 2017). These phenotypic discrepancies may be explained by the possibility that the *dpp* locus is less efficiently excised in *dpp^FO^* flies than in the other two lines (Bosch et al., 2017). In this study we found that the different genetic engineering strategies used to create these conditional alleles likely disrupt the unexcised *dpp* locus to different degrees (Figure 1A and B). In addition to the *FRT* insertion, the *dpp^CA^* and *dpp^PSB^* alleles carry a Hemagglutinin (HA) tagged *dpp* and a visible selection marker (*3xP3-RFP*) in close proximity to the *dpp* locus (Bosch et al., 2017). These insertions might create a weakly sensitized genetic background that explains the different wing growth phenotypes. It is also possible that HA-tagged Dpp is less stable or cannot activate the signaling pathway as efficiently as the *wild-type* Dpp. Indeed, HA-tagging of BMP molecules can result in reduced signaling activity *in vitro* (Figure 5-supplement figure 2) (Akiyama et al., 2012). Therefore, we infer that Dpp protein perdurance and/or residual Dpp expressing cells may be enough to sustain tissue growth until the late third larval instar without the canonical Dpp/BMP activity gradient (Akiyama and Gibson, 2015a). Consistent with this, it has been reported that uniform Dpp expression is able to maintain tissue growth in the wing disc devoid of the stripe Dpp expression, indicating a Dpp/BMP activity gradient independent growth mechanism (Bosch et al., 2017).

### Transcriptional regulation of *nord* in the developing wing

Our transcriptomic analyses using *dpp^FO^* revealed a novel Dpp/BMP target gene *nord* (Figure 1C-I). Surprisingly, unlike other well-studied Dpp/BMP target genes, *nord* expression is restricted in the anterior compartment (Figure 2A-F; Figure 2-figure supplement 1A-H) (Nellen et al., 1996; Tsuneizumi et al., 1997; Campbell and Tomlinson, 1999; Minami et al., 1999; Vuilleumier et al., 2010; Szuperak et al., 2011). We found that Dpp/BMP signaling activity is essential but not sufficient to induce *nord* expression (Figure 2D-I; Figure 2-figure supplement 2A-F). This suggests that other factors are required for its expression. Indeed, a recent study showed that *nord* expression is also regulated by Hh signaling (X. Zheng and MB. O’Connor, personal communication and coordinated submission). Thus, *nord* expression is positively regulated by both Dpp/BMP and Hh inputs, although the molecular basis for *nord* suppression at the D/V boundary remains elusive. Interestingly, *dally* expression is also controlled by these signaling pathways (Fujise et al., 2003). *dally* expression is activated by Hh signaling in the central stripe region of the wing disc, while Dpp/BMP signaling downregulates *dally* expression outside the stripe domain. Thus, through transcriptional regulation, *nord* and *dally* are co-expressed in part of the central stripe region. This may allow Nord to efficiently modulate functions of HSPG Dally to fine tune the Dpp/BMP signaling pathway in the extracellular space.

### Extracellular HSPG regulation by Nord

Extracellular regulation of HSPG, such as shedding of membrane-localized HSPGs from the cell surface, and digestion and/or modification of HS chains, is tightly regulated during development and homeostasis, and its misregulation has been recognized as a hallmark of cancer (Hammond et al., 2014; Nakato and Li, 2016; Xie and Li, 2019; Jayatilleke and Hulett, 2020). Matrix metallopeptidase 9 cleaves a transmembrane form HSPG syndecan-1 from the cell membrane, facilitating tumor growth, metastasis and angiogenesis (Yang et al., 2007; Purushothaman et al., 2008). In addition, the *Drosophila* endo-6-*O*-suflatase Sulf1, which selectively removes 6-*O*-sulfate groups from the HS chains in the extracellular space, controls molecular interactions between HSPGs and signaling molecules including BMP and Wg, and regulates diverse biological processes such as wing development and intestinal stem cell activity (Kleinschmit et al., 2010; Wojcinski et al., 2011; You et al., 2011; Butchar et al., 2012; Dani et al., 2012; Kleinschmit et al., 2013; Takemura and Nakato, 2017). It has also been reported that a secreted Dpp/BMP feedback regulator, Pent, degrades glypicans by endocytosis for the proper formation of Dpp and Wg morphogen gradients (Norman et al., 2016). Here, we showed that Nord regulates Dpp/BMP signaling as an antagonist (Figure 3; Figure 4) and that Nord physically interacts with HSPG Dally to regulate Dpp/BMP signaling in the developing wing (Figure 5A-Q; Figure 5-figure supplement 1). Since Dpp is also a heparin-binding protein and interacts with Dally, we favor a model in which Nord controls Dpp/BMP signaling by modulating co-receptor Dally availability on the cell surface (Figure 5R) (Akiyama et al., 2008). In this framework, Nord competes with Dpp for Dally binding, thus antagonizing Dpp/BMP signaling. Further, a recent study demonstrates a physical interaction between Nord and Dpp (X. Zheng and MB. O’Connor, personal communication and coordinated submission). Therefore, Nord could also regulate Dpp/BMP signaling by directly interacting with Dpp (Figure 5R).

### Implications and conclusions

*nord* encodes a *Drosophila* NDNF protein. Human NDNF acts as a tumor suppressor (Xia et al., 2019; Zhang et al., 2019). NDNF expression is decreased in lung adenocarcinoma, and downregulation of NDNF promotes tumor growth in a mouse xenograft model (Zhang et al., 2019). NDNF is also known as a causative gene for congenital hypogonadotropic hypogonadism (CHH), which is characterized by infertility and delayed/absence of puberty (Messina et al., 2020). CHH is rooted in abnormal development of gonadotropin-releasing hormone (GnRH) neurons. Many CHH-linked genes are involved in the regulation of the Fibroblast growth factor (FGF) signaling pathway (Neocleous et al., 2020). Indeed, NDNF overexpression inhibits FGF signaling in cell culture, and *ndnf* mutant mice exhibit a GnRH neuronal migration defect (Messina et al., 2020). Further, NDNF improves cardiac function after ischemia and myocardial infarction by reducing cardiomyocyte apoptosis and promoting angiogenesis through activation of the AKT signaling pathway (Ohashi et al., 2014; Joki et al., 2015; Song et al., 2017). Nevertheless, the molecular mechanisms by which NDNF modulates multiple signaling pathways remain unclear. Intriguingly, it has been shown that HSPGs play essential roles in these biological processes (Purushothaman et al., 2008; Hammond et al., 2014; Jayatilleke and Hulett, 2020). For instance, mutations in *heparan sulfate 6-O-sulfotransferase-1* and *anosimin-1* are found in CHH patients (Tornberg et al., 2011; Hu et al., 2013; Neocleous et al., 2020). Anosmin-1 is an HSPG interacting protein and regulates GnRH neuronal migration by promoting FGF signaling (Hu et al., 2013). Taken together, we speculate that vertebrate NDNF may modulate HSPG activity to fine tune multiple downstream signaling pathways. In this regard, the present study not only reveals a new mode of extracellular morphogen regulation, but also opens up a new avenue for investigating the molecular basis for NDNF-associated disorders in humans.

## Materials and methods

### *Drosophila* maintenance and stocks

Flies were maintained using standard cornmeal media at 25°C, except where noted. We generated *nord^Δ1^, nord^Δ1728^* and *UAS-nord* as described in detail below. The other fly stocks used in this study were *dpp^FO^* (Akiyama and Gibson, 2015a), *dpp^CA^* (Bosch et al., 2017), *dpp^PSB^* (Bosch et al., 2017), *dpp^hr27^, cn^1^, bw^1^/CyO, P{dpp-P23}RP1* (BDSC#58784) (Spencer et al., 1982; Wharton et al., 1996), *dpp^hr56^, cn^1^, bw^1^/CyO* (BDSC#36528) (Irish and Gelbart, 1987; Wharton et al., 1996), *dpp^FO^; dpp-Gal4/TM6C* (Akiyama and Gibson, 2015a), *dpp^FO^; UAS-FLP* (Akiyama and Gibson, 2015a), *y^1^, vas-Cas9 ZH-2A* (BDSC#66554), *nos-phiC31 int. NLS; attP6* (BDSC#32232), *w^1118^; Df(2R)BSC356/SM6a* (BDSC#24380), *tkv^8^, cn^1^, bw^1^, sp^1^/CyO, P{sevRas1.V12}FK1* (BDSC#34509) (Nusslein-Volhard et al., 1984; Nellen et al., 1994), *Mad^12^, P{neoFRT}40A/CyO, P{GFP-un1}CyO* (BDSC#58785) (Sekelsky et al., 1995), *nub-Gal4* (Calleja et al., 1996), *dally^gem^/TM6B* (Tsuda et al., 1999), *UAS-tkv^QD^* (BDSC#36536) (Nellen et al., 1996), *UAS-Dad* (Tsuneizumi et al., 1997), *UAS-arm^S10^* (BDSC#4782) (Boyle et al., 1997) and *fz3-RFP* (Olson et al., 2011).

### Identification of Dpp/BMP target genes and gene ontology analysis

Wing discs from *Oregon-R* (*wild-type*) and *dpp^FO^/dpp^FO^; dpp-Gal4/UAS-FLP* (Dpp stripeless) larvae were dissected. Total RNA from wing discs was prepared using Direct-zol RNA Miniprep Plus (R2071, Zymo Research). After confirming RNA quality by Bioanalyzer RNA Analysis (Agilent), sequencing libraries were generated using TruSeq RNA Library Prep Kit (Illumina) and sequenced on an Illumina HiSeq system (single read, HiSeq-50bp). 265 genes were differentially expressed in Dpp stripeless wing discs compared to *wild-type* samples (an adjusted *P*-value < 0.01 and a fold change > 1.5). Gene ontology term enrichment analysis of 265 genes was performed on the PANTHER Classification System web site (http://pantherdb.org) (Mi et al., 2019)

### *nord* RNA FISH

1,029-bp *nord* cDNA fragment with T3 promoter was amplified from RE56892 clone (DGRC#9626) using primers 1 and 2 (Supplementary Table) and used for making a Digoxigenin (DIG)-labeled FISH probe using T3 RNA polymerase (P2083, Promega). After T3 RNA polymerase reaction at 37°C for 4 hours, the reaction mixture was treated with RQ1 RNase free DNase (P6101, Promega) at 37°C for 30 minutes to digest template DNAs.

Female third-instar wing discs and 24-hour APF pupal wings were dissected in cold Schneider’s media and fixed with 4% paraformaldehyde (PFA)/phosphate buffered saline (PBS). Wing discs were fixed at room temperature (RT) for 20 minutes and a pupal wing fixation was conducted at 4°C for three days. Fixed samples were dehydrated by 25%, 50% and 75% methanol (MetOH)/PBST (PBS + 0.1% Tween-20) solutions in a stepwise manner. After storing in 100% MetOH at −20°C overnight, samples were treated with 3% hydrogen peroxide/MetOH at RT for 1 hour to eliminate endogenous peroxidase activity and gradually rehydrated using the same series of MetOH/PBST solutions in reverse. Tissue samples were briefly washed three times with PBST and refixed with 4%PFA/PBS at RT for 10 minutes. Samples were equilibrated with 1:1 PBST/Prehybe solution at 37°C for 5 minutes prior to prehybridization at 60°C for 2 hours using Prehybe solution. Samples were hybridized at 60°C overnight in Hybridization solution (Prehybe solution + 0.15 ng/μl *nord* FISH probe). Hybridized samples were washed first with Prehybe and Hybe solutions at 60°C for 30 minutes three times each. Samples were briefly rinsed with PBST, blocked in 1xBM Blocking/PBST at RT for 2 hours and incubated with anti-DIG-POD (1:1,000, 11207733910, Roche) at 4°C overnight. After PBST and TNT washes at RT for 20 minutes three times each, *nord* mRNAs were visualized using TSA Plus Cyanine 5 (NEL745001KT, PerkinElmer). Dpp and GFP expression after *nord* FISH were detected as described below in the Immunohistochemistry and imaging section.

- PBS: 137 mM NaCl, 27mM KCl, 100mM Na_2_HPO_4_ and 18 mM KH_2_PO_4_, pH 7.4
- Prehybe solution: 50% formamide, 5xSSC, 1% SDS, 0.1% Tween-20, 0.05 mg/ml heparin and 0.05 mg/ml salmon sperm
- Hybe solution: 50% formamide, 5xSSC, 1% SDS and 0.1% Tween-20
- TNT: 100 mM Tris-HCl (pH 7.5), 150 mM NaCl and 0.05% Tween 20

### Generation of *nord* mutants

To generate *nord^Δ1^* and *nord^Δ1728^,* we created two sgRNA DNA constructs using primers 3-6 listed in Supplementary Table. Primers were annealed and cloned into the BbsI site of *pBFv-U6.2* (Kondo and Ueda, 2013). A DNA mixture containing two *nord* sgRNA and a *white* sgRNA DNA constructs (100 ng/μl for each) was injected into the posterior region of embryos expressing Cas9 under the *vas* promoter (BDSC#66554). A white eye phenotype was used as a Cas9 activity indicator. Mutant candidates were screened by MiSeq analysis, and the lesions of two mutant lines were reconfirmed by Sanger sequencing.

### *UAS-nord* transgenic line

*nord* coding sequence was amplified from RE56892 clone (DGRC#9626) using primers 7-8 containing an EcoRI site at the 5’ end and an XhoI site at the 3’ end (Supplementary Table). The PCR product was cloned into the EcoRI/XhoI sites of *pUAST attB* (addgene) (Bischof et al., 2007). The resultant *pUAST attB-nord* (200 ng/μl) was injected into the posterior side of *nos-phiC31 int. NLS; attP6* (BDSC#32232).

### Drosophila crosses

Adult viability experiment of *dpp* conditional knockout lines (Figure 1B)

dpp^hr56^, cn^1^, bw^1^/CyO x Oregon-R (control)

-*dpp^hr56^, cn^1^, bw^1^/CyO* x *w^1118.^* (control)

*dpp^hr56^, cn^1^, bw^1^/CyO* x *w; dpp^FO^*

*dpp^hr56^, cn^1^, bw^1^/CyO* x *w; dpp^CA^*

*dpp^hr56^, cn^1^, bw^1^/CyO* x *w; dpp^PSB^*

*dpp^hr27^, cn^1^, bw^1^/CyO, P{dpp-P23}RP1* x *Oregon-R* (control)

*dpp^hr27^, cn^1^, bw^1^/CyO, P{dpp-P23}RP1* x *w^1118^* (control)

*dpp^hr27^, cn^1^, bw^1^/CyO, P{dpp-P23}RP1* x *w; dpp^FO^*

*dpp^hr27^, cn^1^, bw^1^/CyO, P{dpp-P23}RP1* x *w; dpp^CA^*

*dpp^hr27^, cn^1^, bw^1^/CyO, P{dpp-P23}RP1* x *w; dpp^PSB^*

**Dpp stripeless wing disc experiments** (Figure 1E-G, Figure 2G-I)

-*Oregon-R* x *Oregon-R*

*-w^1118^; dpp^FO^; dpp-Gal4/TM6C* x *w^1118^*; *dpp^FO^; UAS-FLP*

**Adult wing phenotype analysis** (Figure 3K)

*w^1118^* x *w^1118^* (control)

*w^1118^; nord ^Δ1728^/CyO, twi-Gal4, UAS-GFP* x *w^1118^; nord ^Δ1728^/CyO, twi-Gal4, UAS-GFP*

*w^1118^; nord ^Δ1^/CyO, twi-Gal4, UAS-GFP* x *w^1118^; nord ^Δ1^/CyO, twi-Gal4, UAS-GFP*

-*w^1118^; nord ^Δ1728^/CyO, twi-Gal4, UAS-GFP* x *w^1118^; Df(2R)BSC356/ CyO, twi-Gal4, UAS-GFP*

-*w^1118^* x *tkv^8^, cn^1^, bw^1^, sp^1^/CyO, twi-Gal4, UAS-GFP*

-*w^1118^; nord ^Δ1728^/CyO, twi-Gal4, UAS-GFP* x *tkv^8^, nord ^Δ1728^/CyO, twi-Gal4, UAS-GFP*

-*w^1118^* x *Mad^12^, P{neoFRT}40A/CyO, twi-Gal4, UAS-GFP*

-*w^1118^; nord ^Δ1728^/CyO, twi-Gal4, UAS-GFP* x *Mad^12^, P{neoFRT}40A, nord ^Δ1728^/CyO, twi-Gal4, UAS-GFP*

*nord* overexpression experiments (Figure 4)

-w^1118^; nub-Gal4/CyO, twi-Gal4, UAS-GFP x w^1118^ (control)

-*w^1118^; nub-Gal4/CyO, twi-Gal4, UAS-GFP x UAS-nord/CyO, twi-Gal4, UAS-GFP*

Genetic interaction between *nord* and *dally* (Figure 5D-Q)

-*w^1118^* x *w^1118^* (control)

-*w^1118^; dally^gem^/TM6B* x *w^1118^; dally^gem^/TM6B*

-*w^1118^; nord ^Δ1728^; dally^gem^/TM6B* x *w^1118^; dally^gem^/TM6B*

-*w^1118^; nord ^Δ1728^; dally^gem^/TM6B* x *w^1118^; nord ^Δ1728^; dally^gem^/TM6B*

**FLP-OUT analysis** (Figure 2-figure supplement 2)

FLP-OUT clones were induced at 37°C for 10 minutes at 48-72 hour after egg laying (AEL).

-*w^1118^, hs-flp; Ay-Gal4, UAS-GFP* x *w^1118^* (control)

-*w^1118^, hs-flp; Ay-Gal4, UAS-GFP* x *UAS-Dad*

-*w^1118^, hs-flp; Ay-Gal4, UAS-GFP* x *UAS-tkv^QD^/TM6B*

-*w^1118^, hs-flp; Ay-Gal4, UAS-GFP* x *UAS-arm^S10^*

***fz3-RFP* reporter assay** (Figure 4-figure supplement 1)

w^1118^; nub-Gal4/CyO, twi-Gal4, UAS-GFP x fz3-RFP (control)

*w^1118^; nub-Gal4/CyO, twi-Gal4, UAS-GFP x UAS-nord, fz3-RFP/CyO, twi-Gal4,UAS-GFP*

### Immunohistochemistry and imaging

Female third-instar wing discs and 24-hour APF pupal wings were used for staining, except for sexually indistinguishable *nord^Δ1728^*; *dally^gem^* animals due to a male larval gonad developmental defect (Figure 5-figure supplement 1D-G). Tissue samples were dissected in PBS, fixed with 4% PFA/PBS (wing disc: RT for 20 minutes, pupal wing: 4°C for two overnight incubations) and incubated with primary antibodies at 4°C overnight. Primary antibodies used in this study were rabbit anti-Dpp^Pro^ (1:100) (Akiyama and Gibson, 2015a), rabbit anti-pSmad3 (1:1,000, EP823Y, abcam) (Akiyama et al., 2012; Akiyama and Gibson, 2015a), mouse anti-Dl (C594.9B,1:500, DSHB) (Bangi and Wharton, 2006; Akiyama and Gibson, 2015a), and chicken anti-GFP (GFP-1010, Aves Labs). Can Get Signal Immunostain Solution B (NKB-601, TOYOBO) was used to dilute primary antibodies. After three PBST (PBS + 0.1% Triton X-100) washes at RT for 20 minutes, samples were incubated with Alexa Fluor conjugated secondary antibodies (1:500 in PBST, Thermo Fisher Scientific) at RT for 3 hours. Samples were washed with PBST as described above and mounted using ProLong Gold Antifade Mountant (P10144, Thermo Fisher Scientific). All images were obtained using a Leica TCS SP5 confocal microscope.

Adult wings were dehydrated using pure ethanol and mounted with Leica Micromount (3801731, Leica). All pictures were taken by a Leica CTR 5000.

### p-Mad and *fz3-RFP* intensity plot profiles and wing size

All wing disc images were collected at the same confocal setting on the same day and analyzed by the RGB profiler of FIJI. The “Measure” function of FIJI was used to analyze sizes of wing discs.

### Cellular fractionation

*pAWH-nord* was generated by the GATEWAY system (Thermo Fisher Scientific). *nord* cDNA amplified from RE56892 clone (DGRC#9626) using primers 9 and 10 was cloned into *pDONR221*, a GATEWAY Entry vector, via the BP reaction (11789020, Thermo Fisher Scientific). After confirming *nord* cDNA sequence in the Entry vector, the cDNA fragment was subcloned from *pDONR221* to *pAWH* (a GATEWAY Destination vector for C-terminal 3xHA tagging, DGRC#1096) using LR Clonase II Enzyme mix (11791020, Thermo Fisher Scientific), resulting in *pAWH-nord*. *Drosophila* S2 cells were transfected with *pAWH-nord*. After 72-hour incubation at 25°C, the cell culture was collected and used as a total fraction. To prepare cell and medium fractions, the culture was centrifuged at 2,000 rpm for 10 minutes at 4°C.

### Heparin-binding assay

Conditioned media from S2 cells transfected with *pAWH-nord* were applied to a 1 mL Hitrap Heparin HP column (17040701, Cytiva). The column was washed with 10 bed volumes of PBS. Heparin-binding proteins were eluted with 1M NaCl/PBS.

### Co-immunoprecipitation

S2 cells were transfected with *pAWH-nord*, *pAW-dally-Myc* and *pAW-dally^ΔGAG^-Myc* in different combinations and cultured at 25°C for 72 hours to allow cells to produce HA and Myc tagged proteins. The cell cultures were solubilized by adding 10% Triton X-100 to a final concentration of 2% and by agitating at 4°C for 1 hour. To prepare Dynabeads-conjugated anti-HA, Dynabeads Protein G (10003D, Thermo Fisher Scientific) and rat anti-HA antibody (3F10, 11867423001, Roche) were mixed and incubated at 4°C for 1 hour with rotation. After centrifugation at 14,000 rpm for 20 minutes at 4°C, the soluble fractions were mixed with Dynabeads-conjugated anti-HA and incubated at 4°C overnight with agitation. Dynabeads-conjugated anti-HA was briefly washed with PBS three times and proteins bound to anti-HA were eluted with SDS sample buffer.

### Western blot

4xSDS sample buffer was used to prepare SDS-PAGE samples. Protein samples were separated by a 4-20% Mini-PROTEAN Precast Protein Gel (4561096, BIO-RAD) and transferred on an Immun-Blot PVDF membrane (162-0177, BIO-RAD) using a Trans-Blot SD Semi-Dry Transfer Cell (1703940, BIO-RAD). The PVDF membrane was blocked for 30 minutes at RT using 5% Blotting-Grade Blocker (170-6404, BIO-RAD)/Tris buffered saline with Tween 20 (TBST) before primary antibody incubation at 4°C overnight. Mouse anti-HA (1:1,000, 16B12, MMS-101P, BioLegend) and mouse anti-Myc (1:1,000, 9E10, 13-2500, Thermo Fisher Scientific) were used as primary antibodies. After three TBST washes at RT for 20 minutes, the membrane was incubated with HRP-conjugated anti-mouse IgG (H+L) secondary antibody (1:10,000, 115-035-003, Jackson ImmunoResearch) at RT for 1 hour. Both primary and secondary antibodies were diluted in 5% Blotting-Grade Blocker/TBST. HA and Myc tagged proteins were visualized on an Amersham Hyperfilm MP (28906845, Cytiva) using SuperSignal WestDuo (34075, Thermo Fisher Scientific).

- 4xSDS sample buffer: 200 mM Tris-HCl (pH 6.8), 8% SDS, 100 mM DTT, 40% glycerol and 0.04% BPB
- TBST: 25 mM Tris-HCl (pH 7.4), 150 mM NaCl, 2 mM KCl and 0.1% Tween-20

### S2 cell-based BMP signaling assay

S2 cell-based signaling assay utilized a *lacZ* reporter whose expression was induced by Notch activation via S(H) binding sites and repressed by BMP silencer elements (Muller et al., 2003; Akiyama et al., 2012). S2 cells were transfected with 10 ng of *pAW-dpp* or *pAW-dpp 3xHA* along with the *lacZ* reporter and other DNAs required for this assay. After 72 hour-incubation at 25°C, S2 cells were lysed to measure β-galactosidase activity using the Dual-Light Luciferase & β-Galactosidase Reporter Gene Assay System (T1003; Thermo Fisher Scientific). Luciferase activity was used for normalizing transfection efficiency.

## Acknowledgements

We thank Hiroshi Nakato for extensive discussion and reagents, Osamu Shimmi for suggestions for pupal wing experiments, Kausik Si for the *white* sgRNA DNA construct, the Bloomington Stock Center for fly stocks, the Drosophila Genomics Resources Center for *nord* cDNA and the Developmental Studies of Hybridoma Bank for anti-Dl antibody. We are grateful to members in the Gibson lab for discussion and advice, especially Lacey R. Ellington for a critical reading of the manuscript and Abby Dreyer for administrative support. We also thank the Stowers Genome Engineering team, Kyle Weaver, MaryEllen Kirkman and Kym Delventhal, for the MiSeq analyses and people in the Stowers Lab Services, Angela Zong, Juan Ramirez and Stacey Walker, for preparing fly foods. This work was supported by funding from the Stowers Institute for Medical Research and NIH R01 GM111733 to M.C.G.

## Author Contribution

T.A. and M.C.G. conceived the project, designed the experiments and wrote the manuscript. T.A. performed the experiments and analyzed the data. C.W.S analyzed the RNA-Seq data.

---

**Figure 2-figure supplement 1:**
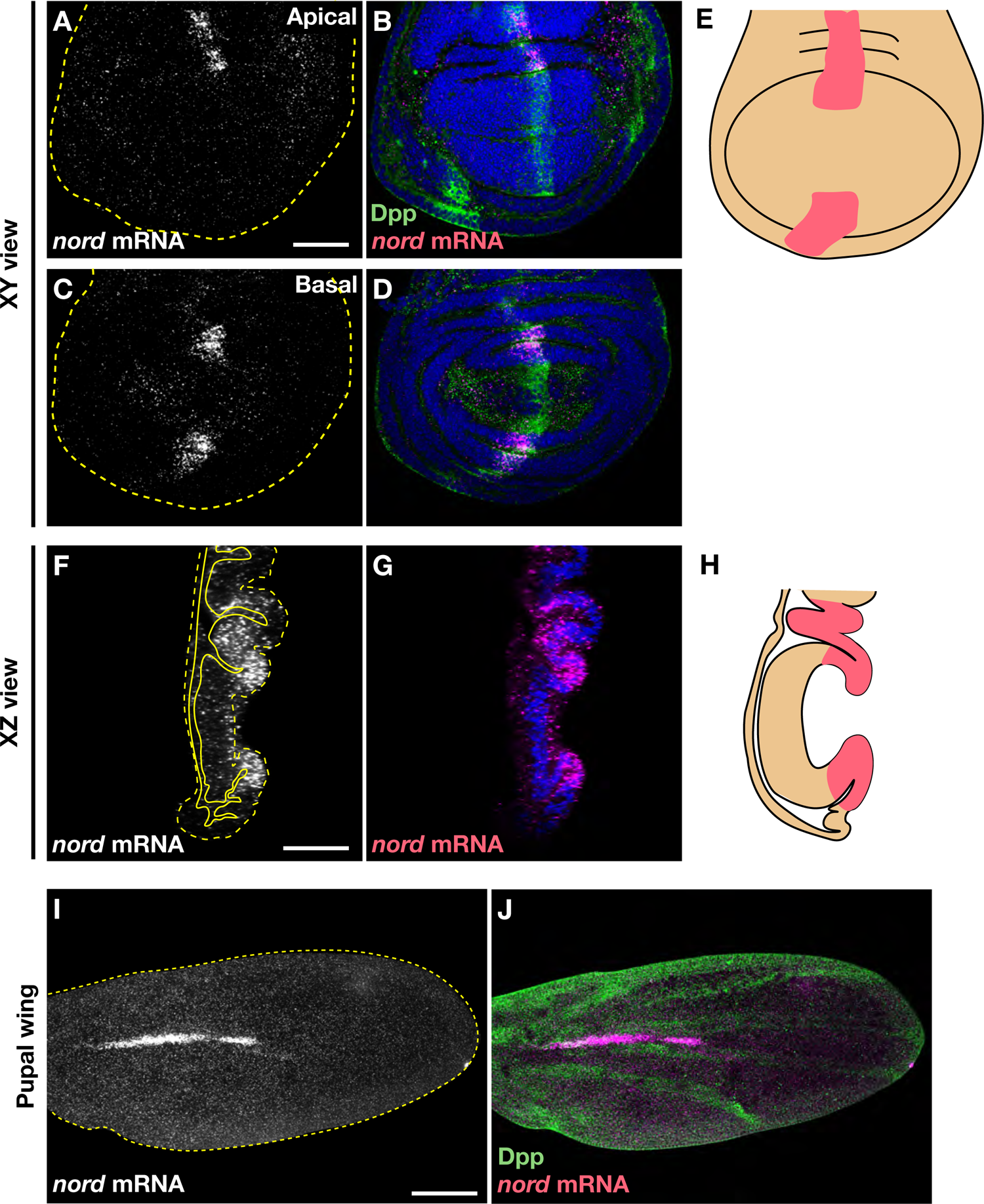
*nord* expression in the wing disc and pupal wing. (**A**-**H**) xy and xz images of *nord* expression in the third-instar wing disc. (**E**, **H**) Illustrations represent *nord* expression in the wing disc. (**I**, **J**) *nord* expression in the 24-hour APF pupal wing. Scale bars: 50 μm for **A** and **F**, 100 μm for **I**. Anterior is oriented to the left in xy wing disc images and to the top in pupal wing panels.

**Figure 2-figure supplement 2:**
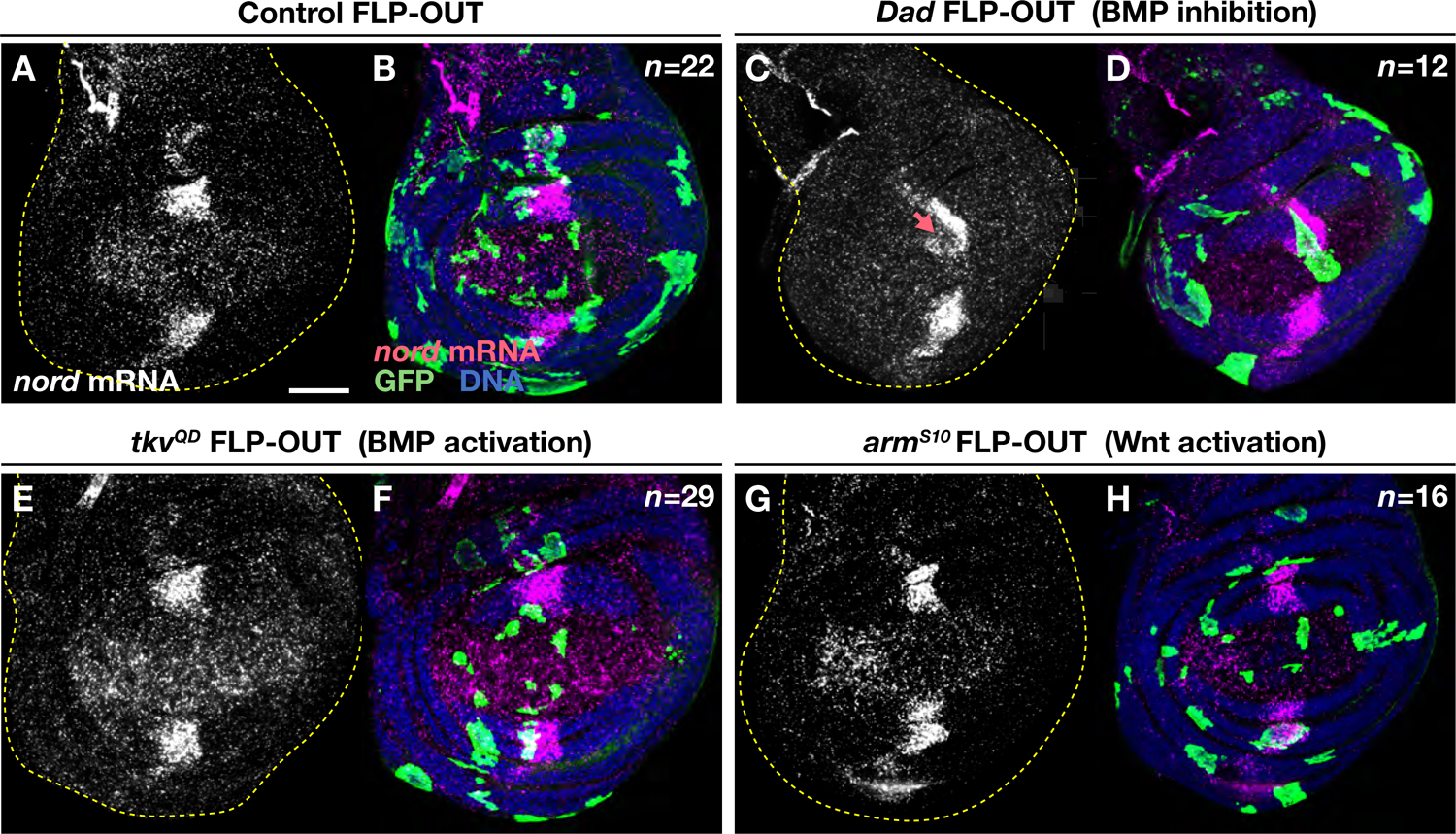
Dpp/BMP is required, but not sufficient, for *nord* expression. (**A**-**H**) *nord* RNA FISH in the wing discs carrying control, *tkv^QD^*, *Dad* and *arm^S10^* overexpressing clones. While *Dad* overexpression strongly reduced *nord* expression (**C**, **D**; arrow), *nord* expression was not clearly affected in *tkv^QD^* and *arm^S10^*clones (**E**-**H**). Scale bar: 50 μm for **A**. Anterior is oriented to the left.

**Figure 3-figure supplement 1:**
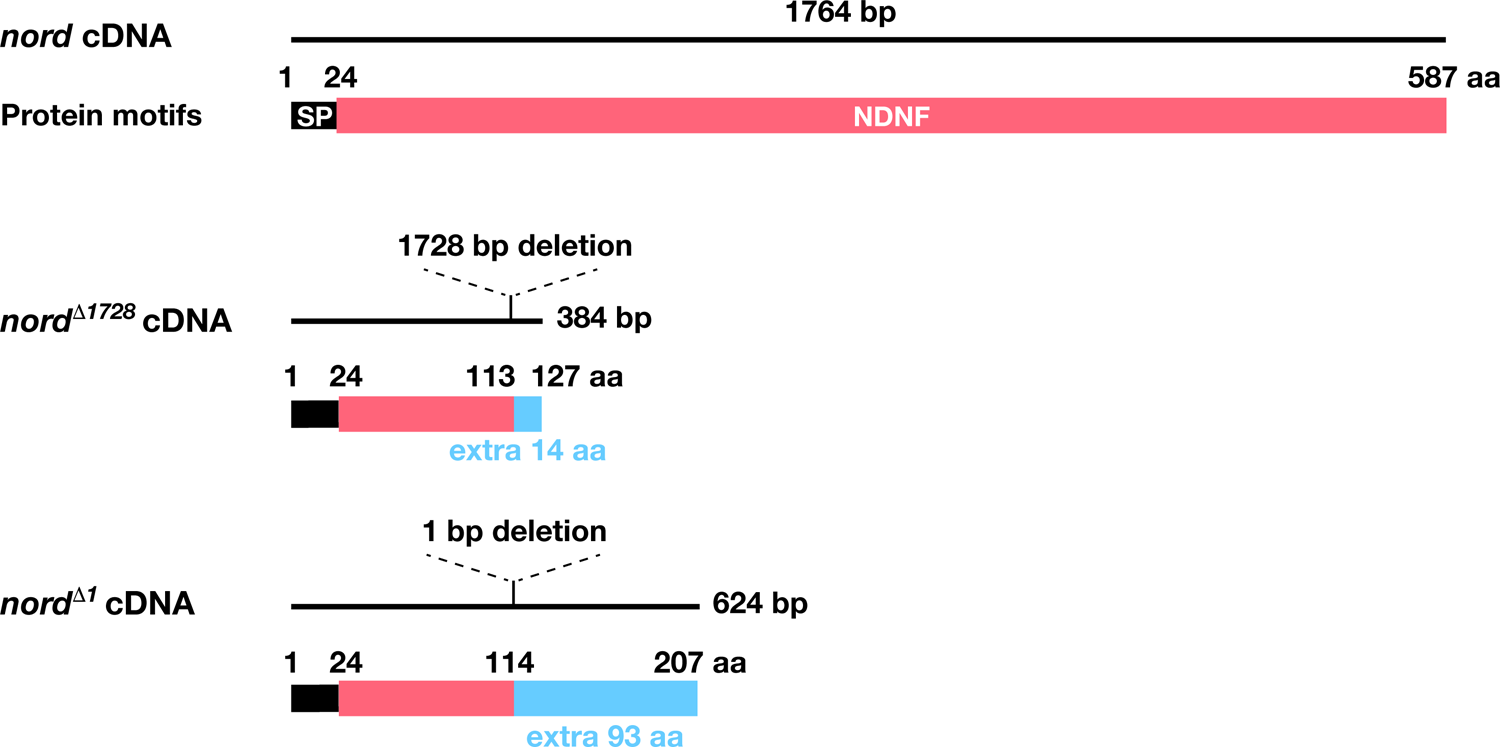
*nord wild-type* and mutant cDNA and protein structures. Lesions *of nord^Δ1728^* and *nord^Δ1^* are mapped on the cDNAs. Both deletion mutations lead to frame shifts, generating protein truncations with extra amino acids (*light blue*). Signal peptide (SP; *black*) and NDNF (*magenta*) motifs are indicated.

**Figure 4-figure supplement 1:**
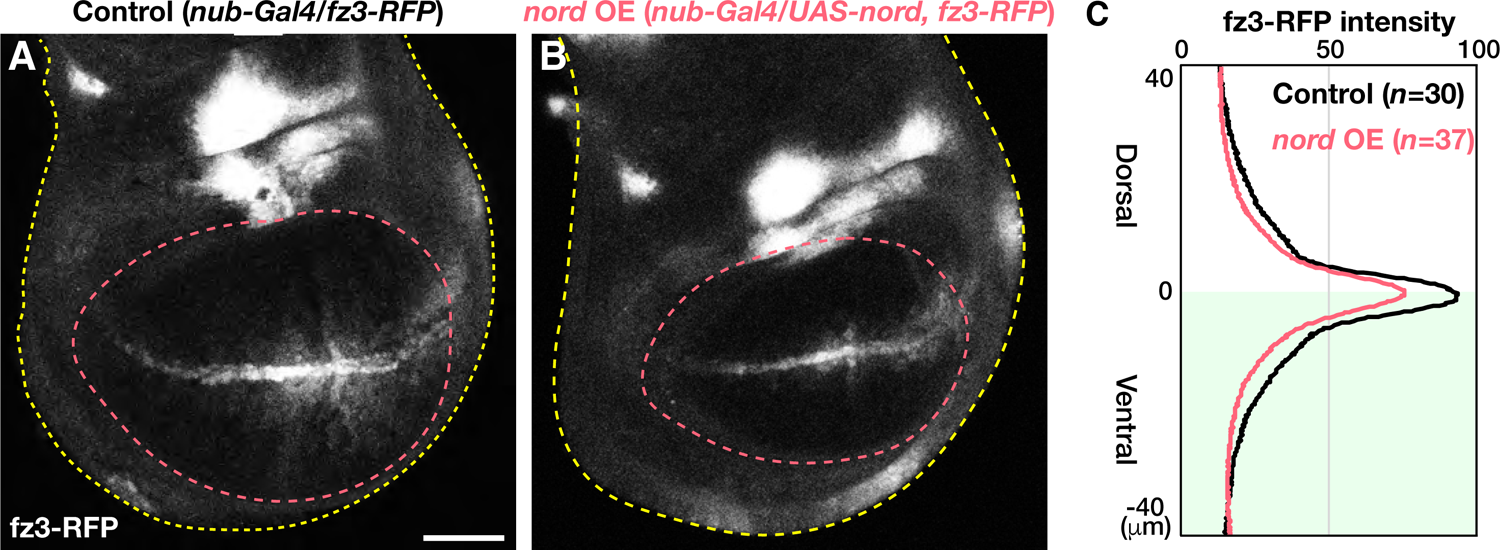
*nord* overexpression downregulates Wg/Wnt signaling activity. (**A**, **B**) Wg/Wnt activities in control and *nord* overexpressing wing discs were visualized by a *fz3-RFP* reporter. *nub-Gal4* induces *nord* expression in the wing pouch (*magenta* dashed line). (**C**) Averaged fz3-RFP intensity plot profiles. *nord* overexpression caused a reduction of fz3-RFP signals. The ventral compartment is shaded. Dorsal side is oriented to the top. Scale bars: 50 μm

**Figure 5-figure supplement 1:**
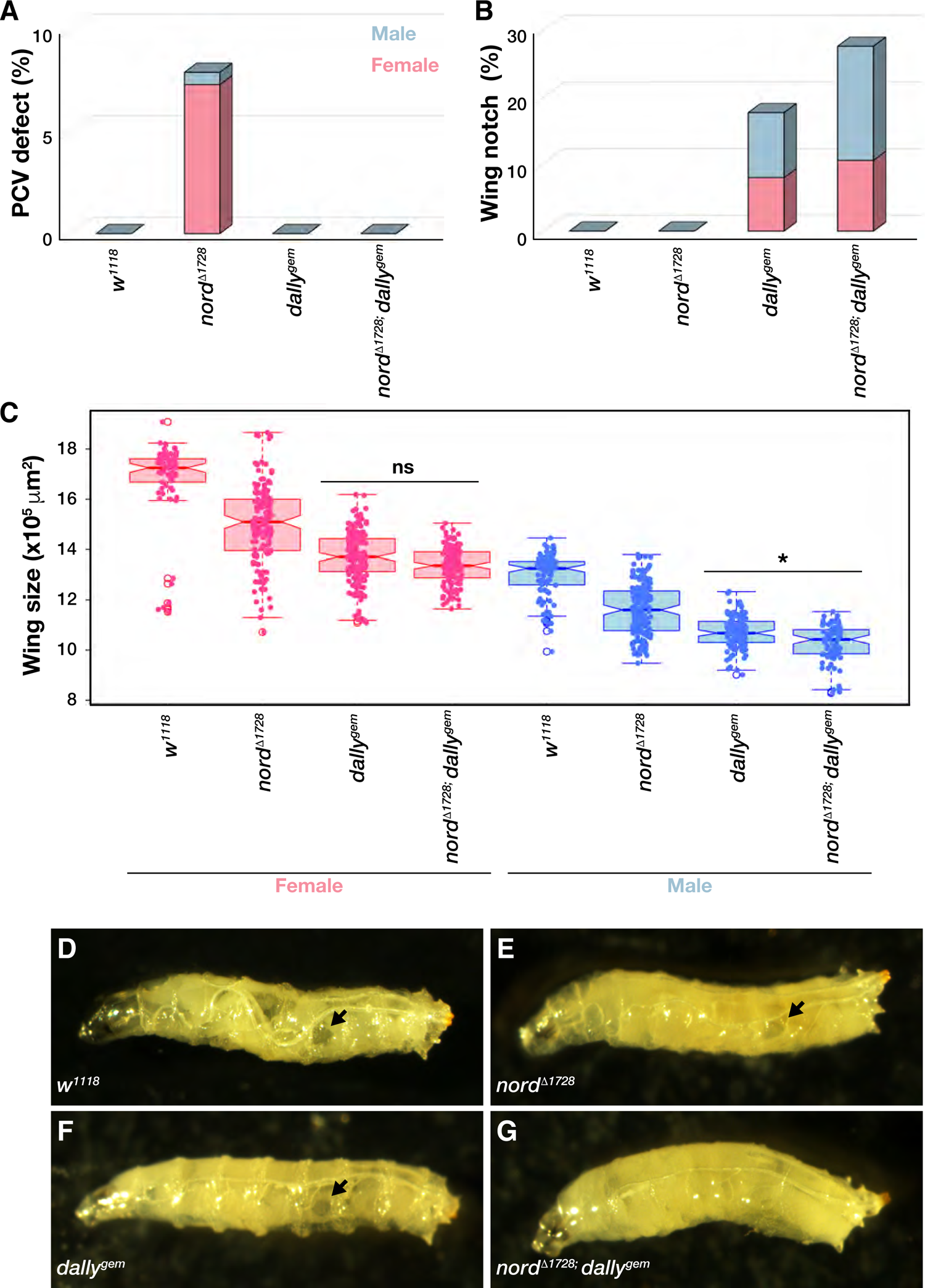
Genetic interaction between *nord* and *dally*. (**A**, **B**) Quantification analyses of the PCV and wing notch phenotypes. The PCV defect was suppressed (**A**), but wing notch phenotype was enhanced in *nord^Δ1728^; dally^gem^* (**B**). *w^1118^* (female: *n*=67, male: *n*=98). *nord^Δ1728^* (female: *n*=150, male: *n*=165). *dally^gem^* (female: *n*=160, male: *n*=117). *nord^Δ1728^; dally^gem^* (female: *n*=146, male: *n*=76). (**C**) Wing size quantification. *w^1118^* (female: *n*=66, male: *n*=96). *nord^Δ1728^* (female: *n*=150; male: *n*=163). *dally^gem^* (female: *n*=153, male: *n*=113). *nord^Δ1728^; dally^gem^* (female: *n*=141, male: *n*=74). \**P*-value < 0.001, not significant (ns), two-sided Student’s *t*-test. (**D**-**G**) A male gonad developmental defect in *nord^Δ1728^; dally^gem^*. The double mutant lacked the male gonad. Arrows indicate circular translucent male gonads in *w^1118^* and the single mutants.

**Figure 5-figure supplement 2:**
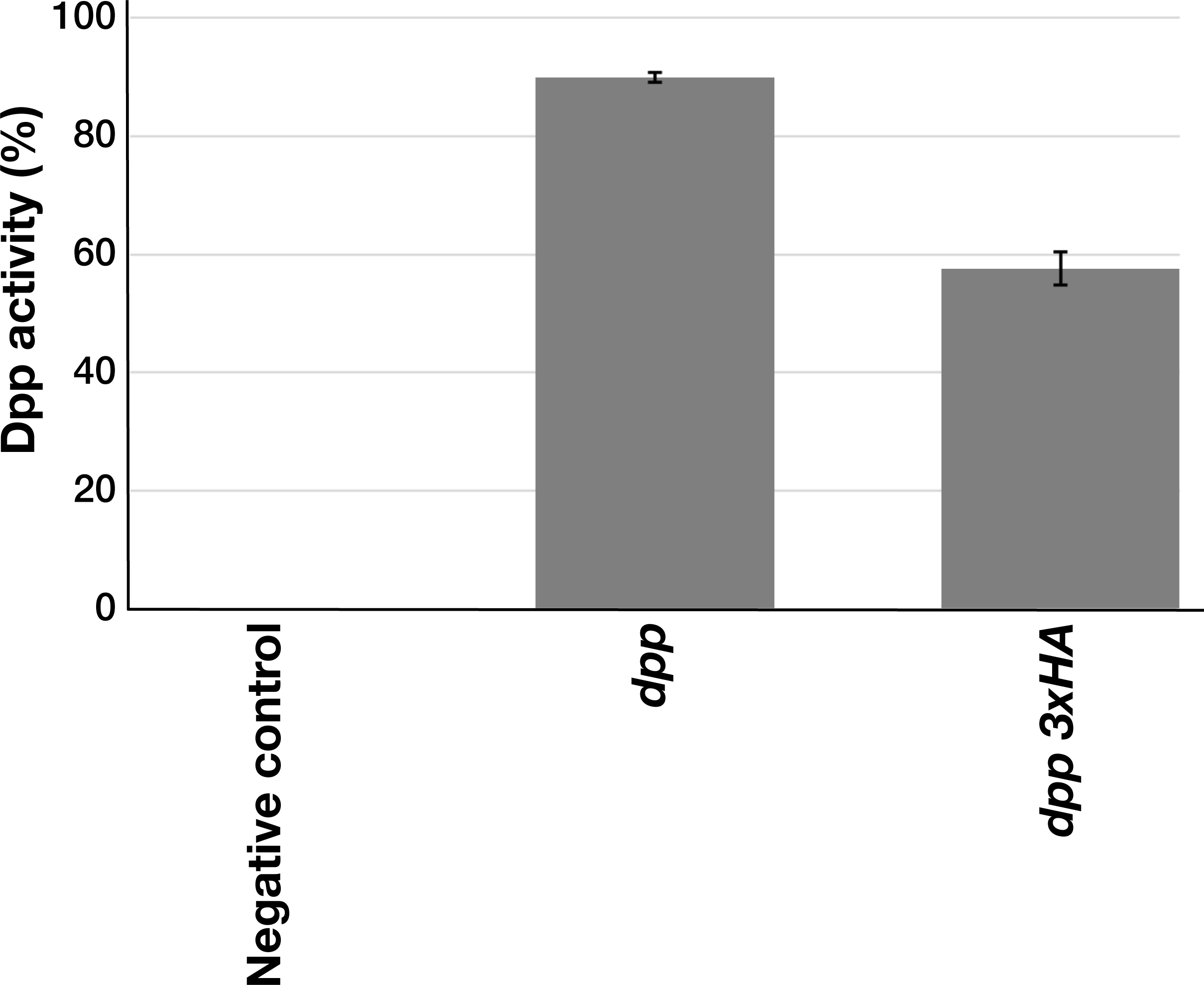
*In vitro* signaling activity of Dpp and Dpp 3xHA. Signaling activity of *wild-type* Dpp and Dpp 3xHA were examined using an S2 cell-based BMP signaling assay. Dpp 3xHA showed a reduced signaling activity compared to Dpp.

**Supplementary Table.**
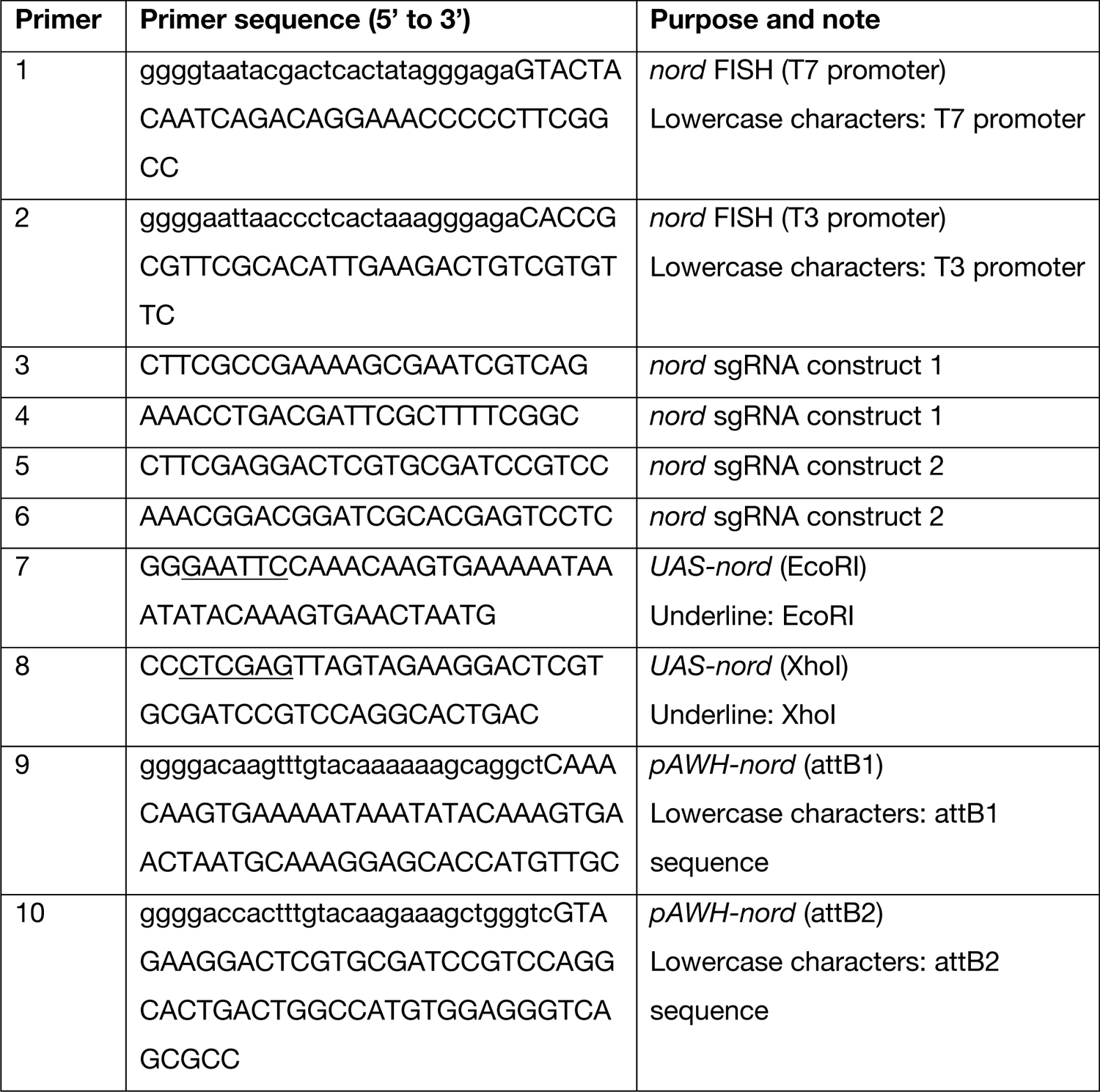
Primers used in this study

